# Bayesian phylodynamics of early vertebrate development in BEAST 2

**DOI:** 10.1101/2024.07.04.601658

**Authors:** Antoine Zwaans, Sophie Seidel, Marc Manceau, Tanja Stadler

**Affiliations:** Department of Biosystems Science and Engineering, ETH Zürich, Basel, Switzerland; Swiss Institute of Bioinformatics, Lausanne, Switzerland

## Abstract

Analysing single-cell lineage relationships of an organism is crucial towards understanding the fundamental cellular dynamics that drive development. CRISPR-based dynamic lineage tracing relies on recent advances in genome editing and sequencing technologies to generate inheritable, evolving genetic barcode sequences which enable reconstruction of such cell lineage trees, also referred to as phylogenetic trees. Recent work generated custom computational strategies to produce robust tree estimates from such data. We further capitalise on these advancements and introduce GABI (GESTALT Analysis using Bayesian Inference), which extends the analysis of GESTALT (Genome Editing of Synthetic Target Arrays for Lineage Tracing) data to a fully integrated Bayesian phylogenetic inference framework in the software BEAST 2. This implementation allows to represent the uncertainty in reconstructed tree reconstruction and enables their scaling in absolute time. Furthermore, based on such time-scaled lineage trees, the underlying processes of growth, differentiation and apoptosis are quantified through so-called phylodynamic inference, typically relying on a birthdeath or coalescent model. After validating the implementation, we demonstrate that the methodology results in robust estimates of lineage trees and growth dynamics characteristics of early zebrafish *Danio rerio* development. GABI’s codebase is publicly available at https://github.com/azwaans/GABI.

## Background

The underlying principles of multicellular growth remain a major question in biology. Understanding the dynamics of cell division is central to the study of biomedical processes ranging from healthy embryonic development, organogenesis and tissue regeneration to cancer progression and tumour growth [1]. In fact, both healthy and diseased tissues are fundamentally the result of a complex balance between the process of cell proliferation and death. Considerable efforts have been made towards gaining insights into the timing, rate, and modality of these processes. Their complexity lies in the fact that cell fate is determined through the spatial and temporal integration of a plethora of factors, that include endogenous pathways, long distance hormonal or even biophysical signalling, to only name a few [2]. This is partly addressed by a growing body of experimental methods producing genomic, transcriptomic and proteomic data at the single cell level [3]. These increasingly comprehensive, multiomics datasets provide the resolution required to characterise individual cell states [4]. However, this molecular resolution of cells at particular time points does not give direct information on the changes of the cells through time.

A natural representation of cell populations through time are lineage trees, also referred to phylogenetic trees, where a node depicts a sampled cell, and edges represent ancestry of these cells, with their lengths corresponding to calendar time. Phylogenetic trees can generally be employed to estimate parameters of the processes generating these trees; such inference of population dynamic parameters based on phylogenetic trees is also called phylodynamic inference. Starting with Yule 100 years ago [5], seminal papers demonstrated that birth and death rates in a population can be estimated from a tree with individuals sampled from the present day population [6, 7]. Translated to our single-cell application, we can estimate past division (birth) and apoptosis (death) rates or sampling intensity based on phylogenetic trees reconstructed from single cells sampled at one time point of development [1]. The concept of observing cell lineage trees under the microscope dates back to the late 19th century [8]. However, microscropic observation of such lineages quickly becoming a challenge for large, non-transparent organisms, prompted the usage of sequencing data as a powerful alternative to encode relatedness between cells. While for cancer cells, enough changes in the genome may occur to be able to reconstruct lineage trees [9, 10], typically somatic mutations in healthy cells are too rare to give lineage signal at a high resolution [11]. Instead, drawing on recent developments in sequencing and genome engineering technologies, cell lineage tree reconstruction now increasingly relies on sequences derived from artificial genetic constructs called *lineage recorders* that dynamically accumulate mutations during development. These methods, collectively termed *dynamic lineage tracing* [12] require generating transgenic organisms harbouring the genetic components of the system of choice and use various gene editing strategies to generate mutations. Amongst others [13–15], multiple methods have been developed using the growing toolbox of CRISPR Cas-9 genome editing based technologies. There, the endogenous, error-prone DNA repair mechanisms of double stranded breaks result in the stochastic insertion or deletion of sequences. GESTALT (**G**enome **E**diting of **S**ynthetic **T**arget **A**rrays for **L**ineage **T**racing, [16]), ScarTrace [17] and LINNAEUS (LINeage tracing by Nuclease-Activated Editing of Ubiquitous Sequences [18]) methods all rely on this principle. These repair outcomes constitute the building blocks of sequencing data generated with such methods.

The fields of macroevolution and genomic epidemiology rely on a variety of long established, computational methodologies for the reconstruction of phylogenies from genetic sequencing data [19–23]. However, lineage recording data break traditional assumptions of these methods, in particular regarding molecular evolution [24]. Early examples of lineage trees constructed from genetic lineage tracing data have relied on generic methods such as neighbor-joining [25] or Camin-Sokal parsimony [26] for tree reconstruction, both of which do not model the exact mutation process of lineage tracing barcodes. This can result in poor accuracy in tree reconstruction as these fail to incorporate unique features of the data, such as indel states or the irreversibility of the recording process [27]. This led to recent methodological developments that more closely account for specific features of the data [28], including machine learning methods [29], and custom distance or parsimony based methods [30, 31]. While some of these methods were shown to greatly improve the accuracy of lineage tree reconstruction, they focused exclusively on reconstructing phylograms, i.e. lineage trees that are not scaled in absolute time. Time-scaled lineage trees, or chronograms, additionally hold information on the timing of lineage segregation events and, further, the underlying population dynamics of the studied cell populations [32]. Towards that goal, generating such time-scaled trees was made possible with the formulation of mechanistic models of lineage recording data, used in a Bayesian or Maximum Likelihood inference framework [33–35]. In a previous paper [33], we developed a Bayesian phylodynamic method (Tide-Tree) for single-cell lineage tracing datasets, enabling the joint estimation of lineage trees and cell population dynamics parameters, with an analysis of mouse embryonic stem cells [36]. For GESTALT data specifically, a first mechanistic model describing detailed features of its barcode mutation process was introduced in [37] where cuts introduced by Cas-9 upon recognition of the target sequences are assumed to follow a continuous time Markov chain process. The implementation of the phylogenetic likelihood under that model for maximum likelihood inference, GAPML (GESTALT analysis using penalized Maximum Likelihood), reconstructed lineage trees with a relative timing of division times under a molecular clock assumption. This implementation however provided no scaling of branch lengths in absolute time and no straightforward way to assess uncertainty in estimates.

In this work, we introduce GABI (GESTALT analysis using Bayesian Inference), which implements and extends the likelihood described in [37] into the Bayesian inference software BEAST 2 [20] for joint estimation of lineage trees scaled in absolute time and cell population dynamics from GESTALT sequencing data. We couple this method with the birth-death and coalescent models for tree generation [7, 38]. Both models enable the posterior estimation of the rate of cell divisions and the proportion of cells being sequenced. We validate this framework, and demonstrate its use with an empirical study of *Danio rerio* early development from alignments of GESTALT sequences collected from dome stage embryos. These analyses show that timescaled embryonic lineage trees and the growth rates that underlie this embryonic stage can robustly be inferred using our new framework.

## Methods

### Mutation model

We designed GABI to consider GESTALT barcode alignments as data for Bayesian phylogenetic inference. We enable this by the implementation of a phylogenetic likelihood (i.e. the probability of the sequence alignment given the tree) under the model of GESTALT barcode evolution described in [37] (called GAPML model), in BEAST 2 [20]. In the following, we briefly introduce the notations, definitions and assumptions that were used to formulate this model of GESTALT sequence evolution and underline key assumptions and approximations used for the likelihood calculation under that model.

### GESTALT barcode

GESTALT relies on a Cas9 based technology to generate stochastic mutations introduced at specific sites within a transgene (referred to as barcode) built into the founding cell of the population whose lineage is traced. Lineage data consists of an alignment of such modified transgenes, generated through sequencing of GESTALT barcodes after cell populations of interest undergo proliferation. The following are general notations used to describe any of the several GESTALT barcode designs introduced in [16] but also hold for all subsequent GESTALT applications and related technologies [39–44], and can be directly modelled by only adjusting the available parameters.

The GESTALT barcode is a DNA sequence that consists of an array of *M* subsequences each containing a CRISPR-Cas9 target site. Each of these subsequences contains a protospacer adjacent motif (PAM) sequence necessary for Cas9 nuclease binding and a spacer sequence complementary to single-guide RNA (sgRNA) for specific recognition. In the presence of the Cas9 enzyme, and a sgRNA complementary to the spacer sequence at a target site, mutations of GESTALT barcode sequences occur as the result of a 2-step process: (i) upon sgRNA binding at a target site *k*, Cas9 first generates a double strand cut at a position 3 bp upstream of PAM sequences, denoted as *c*(*k*), and (ii) this then triggers an endogenous repair mechanism that results in the introduction of insertions and deletions into the barcode at the targeted sites (see Appendix Fig. 1, upper half). Introducing an insertion or deletion at each of the *M* target sites is irreversible as it breaks complementarity with its corresponding sgRNA. Therefore, such sets of insertions and/or deletions (together, referred to as *indels*), are the building blocks of characters in GESTALT sequence alignments. For such GESTALT sequencing data, [37] suggested a model of this evolution, implemented in GAPML. It contains parameters that specify target specific cut rates, weights for the relative frequency at which double cut events occur, and factors controlling the frequency of long and short deletions left and right of these cuts (see Table 1 for notations). In Appendix B, we describe the precise model assumptions and how the likelihood is calculated under that model. In our implementation, we additionally introduce a clock rate *r*, quantifying the rate per time unit of mutations per target site (see also Table 1). In GAPML, for any branch of relative length *τ*_*i*_ relative to total tree height, transition probabilities are expressed as elements of the transition probability matrix 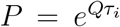. In GABI, we additionally provide scaling in absolute time with the clock rate, where a branch *i* accumulates indels at a rate of *r*_*i*_ indels per target site per unit of time. In BEAST 2, these branch rates can then either be specified as being constant for all branches, corresponding to a *strict molecular clock* ([45])assumption, or allowed to vary across branches or clades, corresponding to a *relaxed clock* ([46][47]). Transition probabilities are obtained as described in Appendix B, but substituting for the absolute branch length *t*_*i*_ instead 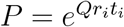.

**Table 1:** Glossary: Parameters of the GABI editing model. All parameters but *r* are assumed in the GAPML method, *r* is additionally introduced here.

| GESTALT parameter | Description |
| --- | --- |
| Molecular clock rate $r$ | Rate at which the barcode acquires mutations, in indels per site per time unit |
| Cut rate $\lambda_i$ | Rate at which an individual target $i$ in the barcode gets cut |
| Double-cut weight $\omega$ | Weight that specifies how frequent double-cut events are relative to single cut events |
| Long-trim factor $\gamma_0$ | Factor specifying how frequent long deletions are relative to short deletions, left of a cut |
| Long-trim factor $\gamma_1$ | Factor specifying how frequent long deletions are relative to short deletions, right of a cut |

### Implementation

We implemented the phylogenetic likelihood and the sequence simulation engine for the GESTALT model from [37] in Java for the Bayesian phylogenetic inference platform BEAST 2, as a package called GABI (GESTALT Analysis using Bayesian Inference). This implementation enables its interaction with dozens of tree priors, molecular clock models, and a growing body of tutorials to facilitate the specification of complex analyses. Its integration within BEAST 2 also ensures interoperability with software that provide visualization tools to summarise MCMC posterior samples and assess convergence.

Changes made from the original Python framework with regards to the tree inference are the following: GABI considers exclusively bifurcating trees and thus does not implement the scheme aimed at resolving multifurcations by creating spines, and does not restrict the tree search to maximum parsimony topologies. Additionally, an advancement made in GABI is the speed-up of the likelihood calculation by caching partial likelihoods and ancestral states for unchanged parts of the lineage tree, leading to up to a 2.5 times speed-up over 100 iterations of the MCMC chain. Further speed-up can be achieved by the use of threading natively supported by BEAST 2.

As outlined above, the core novel component of GABI is its ability to specify and infer a molecular clock rate, *r*, which denotes the rate that converts branch lengths into calendar time. This can further be extended by specifying branch or clade specific rates, when deemed appropriate. Building up on this component, our implementation of a sequence simulation engine, based on BEAST 2 package feast [48], is the first fully integrated simulator allowing to generate GESTALT alignments along time-scaled lineage trees, while specifying branch specific mutation rates. Beyond assessment of proper implementation as shown here, it also provides a framework for more investigations of the properties of GESTALT data with respect to its ability to encode signal for both lineage, and cellular growth patterns.

Indel sequences characteristic of GESTALT data are not supported natively as input for BEAST 2. To enable their use, we defined a new BEAST 2 datatype format, which requires indels to be encoded with their start position in the barcode, the deletion length, the target sites affected by the deletion and the insert sequence in this order. GABI also enables to specify any GESTALT barcode design of interest as metadata associated with input GESTALT alignments, containing the unedited barcode sequence and the position of the cut site relative to the 3’ end of each target site. All helper functions that are part of the GABI package interact as shown in Appendix Figure 2.

### In-silico validation

We first validate the correct implementation of the phylogenetic likelihood by comparing its output values with the previous Python implementation of GAPML. For this, we calculate the likelihood output for 13 datasets that cover all mutation types (unedited barcode, focal deletion or long deletions) and a range of phylogenetic structures (single branch, 3-leaf tree and 4-leaf tree). For each of these tests, we additionally record the runtime needed for a single, initial evaluation of the likelihood in both implementation, recorded on a computer running OSX on Intel(R) Core(TM) i5 Dual-Core @2.30Ghz.

We examine the statistical correctness of the likelihood using well-calibrated simulations [49]. We first draw 100 sets of mutation parameters from distributions described in Appendix Table 2. For every mutation parameter set, we simulate a dataset along one of 100 simulated trees chosen to have between 50 to 100 tips (and simulated under parameters described in Appendix Table 1). Distributions from which the parameter sets were drawn were centered around previously inferred maximum likelihood values on V7 data of the GESTALT barcode [37]. We chose to simulate alignments under a mean molecular clock rate of ∼ 0.02 indels per site per hour, and arbitrarily assume an experimental duration of 25 days. Together, this is consistent with a relatively low mutation rate to reflect a lower bound in recording capacity, corresponding to an average of about 0.5 indels per barcode sequence in each simulated GESTALT alignment. For simplicity (but without loss of generality), we choose an unedited barcode identical to version 7 (V7) of the GESTALT design [16], but shortened to 4 positions instead of 10, for simplified interpretation. Using these simulated alignments as data, we infer the mutation parameters with the priors being equal to the distributions which were used for simulation, fixing the tree to its true value. Each dataset was used for inference using MCMC for 10^6^ iterations and a burn-in of 30% of the iterations was discarded. Convergence was defined by reaching an expected sample size (ESS) ¿200 for all parameters. Coverage at a given credibility level *α* is calculated for each of the parameters as the proportion of times where the true parameter is contained within the inferred *α* % Highest Posterior Density (HPD) interval over all simulated datasets.

### Phylodynamic analysis of dome stage zebrafish Danio rerio datasets

We showcase GABI’s implementation with a phylodynamic analysis of the growth of dome stage transgenic *Danio rerio* embryos, from [16]. In this experiment, zebrafish embryos are harvested for sequencing of GESTALT barcodes 4.3 hours post fertilisation (hpf) (also see Fig. 1, A). This is a well characterized developmental stage, where embryos are known to have undergone between 12 and 14 rounds of cell divisions and no cell death [50]. This makes the dataset ideal as a first, proof-of-concept application of our framework for the study of cell growth dynamics using GESTALT barcode data. Based on the sequencing data, we aim to infer the time scaled phylogenetic trees together with the division rates (birth rates) and sampling intensity, employing GABI together with the birth-death and coalescent framework in BEAST 2 [7, 20, 38].

**Figure 1:**
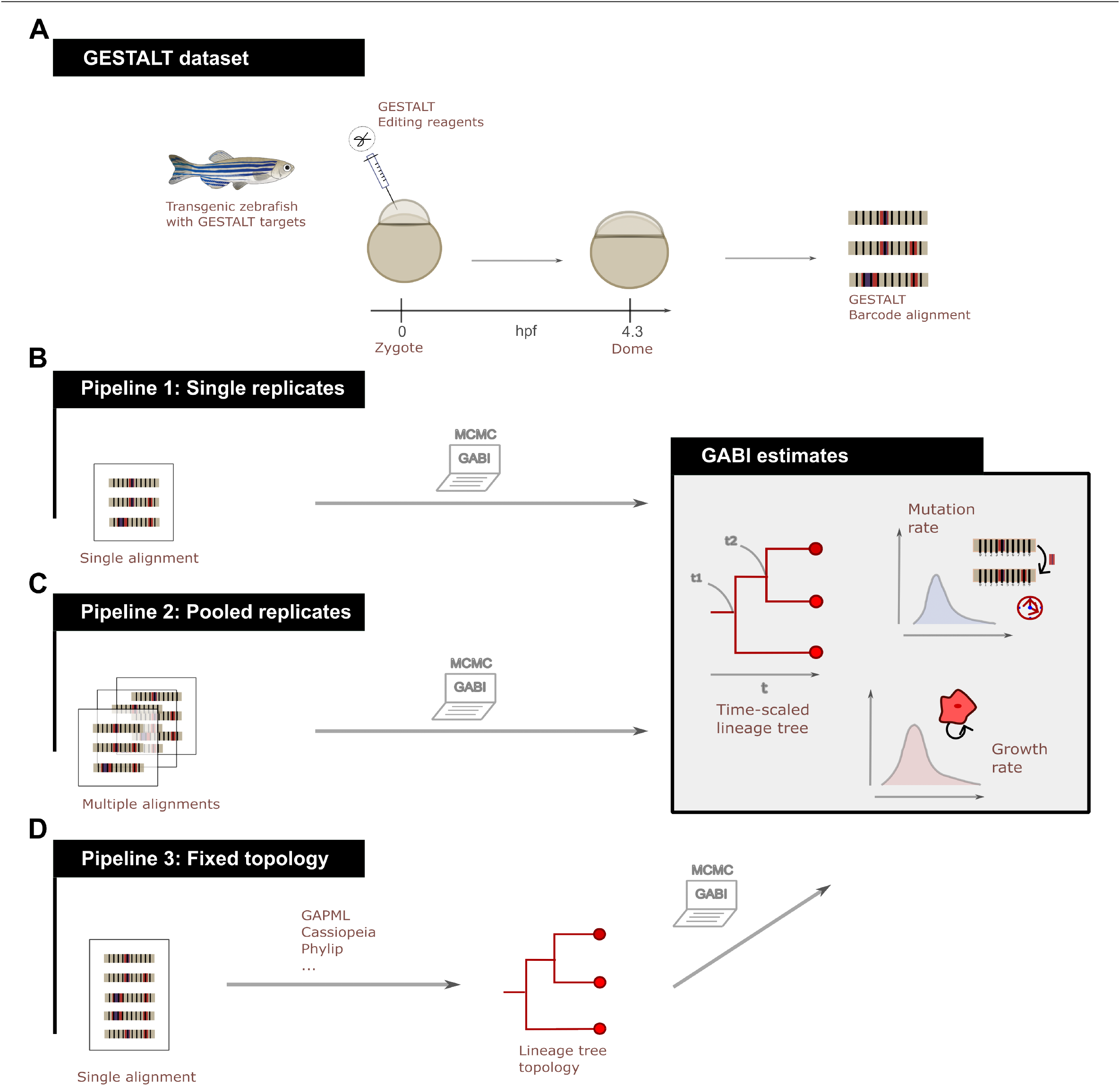
Phylodynamic inference pipelines using GABI. Using GESTALT data obtained for dome stage (4.33 hpf) transgenic zebrafish *Danio Rerio* (A), we showcase the use of GABI using three realistic pipelines, analysing three replicates of the dataset. (B) Pipeline 1: GABI can be applied for inference on individual experimental replicates, inferring a tree topology, branch lengths, as well as mutation and growth parameters. (C) Pipeline 2: when available, replicates of a same experiment can be pooled to obtain more confident estimates. (D) One can also fix a tree topology to one inferred using other methods, and apply GABI to time-scale this lineage tree and obtain parameter estimates.

The GESTALT barcode design used for this experiment is V7, consisting of 10 target sites. Editing of these barcodes is achieved by injection of Cas9 editing reagents into transgenic fertilized *Danio Rerio* eggs, and is thought to taper starting dome-stage [16]. To model these editing dynamics, we assume a constant, strict molecular clock for the duration of the experiment. Prior distributions on the GESTALT editing parameters around mean values estimated previously on the same barcode design (see Appendix Table 3 also, in-silico validation). As estimates of the mean indel lengths were shown to be most consistent over different experiments using barcode V7 in [37], we fix them in this example. This dataset consisted of 10 replicates with a median of 2000 sequences available per embryo. Thus, it constitutes an ideal case study for a first application of phylodynamics to the quantification of zebrafish embryonic development. To model embryonic growth and the harvesting process, we apply the 2 foundational models of population growth available for phylodynamic inference: the birth-death-sampling model [51], and the coalescent with exponential growth [38]. In the birth-death-sampling model, birth rate *β* corresponds to the rate of cell divisions while death rate *δ* (in our applications, fixed to 0) corresponds to the rate at which cells undergo apoptosis, and *ρ* corresponds to the fraction of cells used in the analysis over all cells at the end of the experiment. Under this model, the expected population size after time *t* is *N* (*t*) = *e*^(*β*−*δ*)*t*^.

The coalescent with exponential growth is parameterised with a population growth rate *g* also corresponding here to the rate of cell divisions (assuming no apoptosis happens). Here, the model assumes a population size *N* (*t*) = *e*^*gt*^ at time *t*. Under both models, we report estimates of the cell division rate, and the sampling proportion. For the coalescent analyses on an alignment of *s* cells, the sampling proportion is reported as the ratio 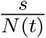 (see Appendix Tables 4 and 5 for detailed priors).

To present different use cases of GABI, we designed 3 analysis pipelines for this dataset, described in Figure 1 B-D. In the first one, we infer the lineage tree topology, its branch lengths, and growth parameters for 3 of the dome alignments individually (Fig. 1 A). For this analysis, we limit the number of sequences per dataset to 20 sequences each to remain within short runtime. As these datasets are replicates of the same experiment, we additionally perform a second analysis where parameters of the GESTALT barcode and cell population growth parameters are inferred over all 60 sequences pooled together (Fig. 1, C). The third analysis pipeline uses a larger subsample of 100 sequences for the first replicate, where the tree topology is fixed to a maximum-likelihood estimate obtained with GAPML (Fig. 1, C). For this analysis, only branch lengths in the lineage tree are estimated jointly with model parameters. Each of these analyses was run for ∼ 10^7^ iterations and ∼ 20% of steps were discarded as burn-in. Convergence is confirmed for parameter estimates that reach an effective sample size (ESS) larger than 200 (and assessed using software Tracer [52]). All trees shown in this study are maximum clade credibility (MCC) trees constructed to summarize the posterior tree sample.

All datasets analysed were obtained from the Gene Expression Omnibus repository, with query GSE81713. Specifically, the 3 datasets analysed are the following: GSM2171788 (dome 1) GSM2171791 (dome 2) GSM2171791 (dome 3), where the provided GESTALT sequences were aligned and processed and aligned as pipeline described in [16]. Briefly, after multiple sequence alignment with the unedited barcode using the MAFFT [53] software, mutations are identified as indels occurring within 3 bp of the predicted cut sites. Additional processing of ambiguous indels is performed using the pipeline provided in [37], and produces the input data for our implementation.

## Results

### In-silico validation

To show correct implementation of the GESTALT phylogenetic likelihood computed in GABI, we first ensured that its output coincides with its previous implementation in Python. GABI implements a special case of the Python implementation of GAPML, where only bifurcating trees are considered. Correct calculation of transition probabilities alone is first assessed by calculating likelihoods along single branches, for all possible indel types. The full likelihood calculation, that recursively calculates and combines subtree likelihoods using Felsenstein’s pruning algorithm is then tested on 3 small bifurcating trees with example sequences. The Figure 2 A) shows equality of the log-likelihood values calculated under both implementations. Additionally, we recorded the runtime needed to calculate these likelihood values under each implementation (see Fig. 2 B) and find that GABI outperforms the original Python code, by up to a order of ∼ 100 fold (which corresponds to up to seconds).

**Figure 2:**
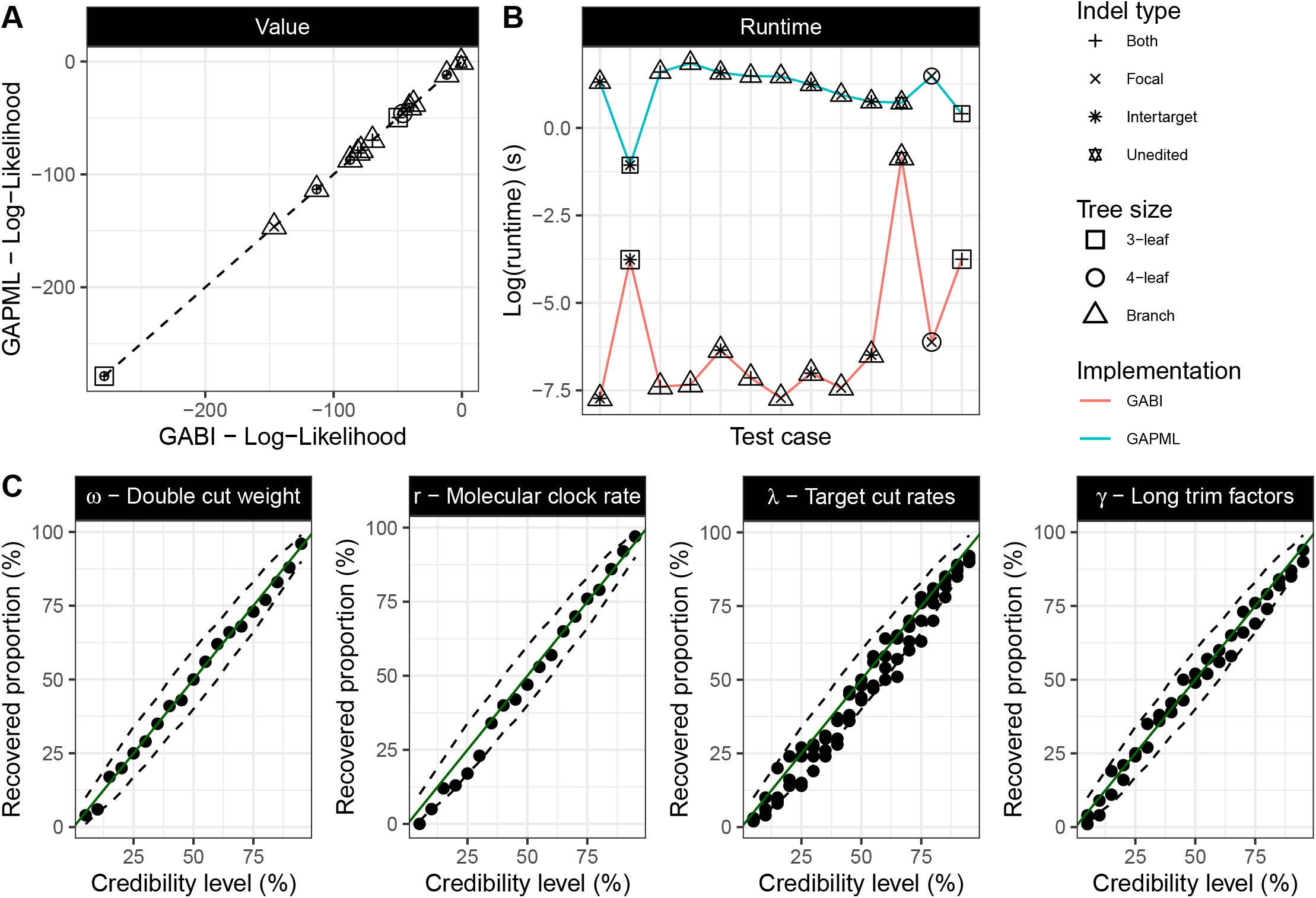
In-silico validation. (A) Indicates the likelihood implementation correctness with a direct comparison of the likelihood values calculated in GABI and in GAPML for different test cases (see legend). The dotted line shows the expected equality between both values. (B) Shows the log(runtime) evaluated on single, startup iterations of the likelihood calculation under both implementations for the same test cases. (C) Indicates the MCMC framework correctness. Each dot represents the coverage obtained for credibility levels between 0.05 and 0.95. Dashed lines represent the 95% Confidence Interval of a Binomial distribution with *n* = 100, and probability of success between 0.05 and 0.95.

**Figure 2:**
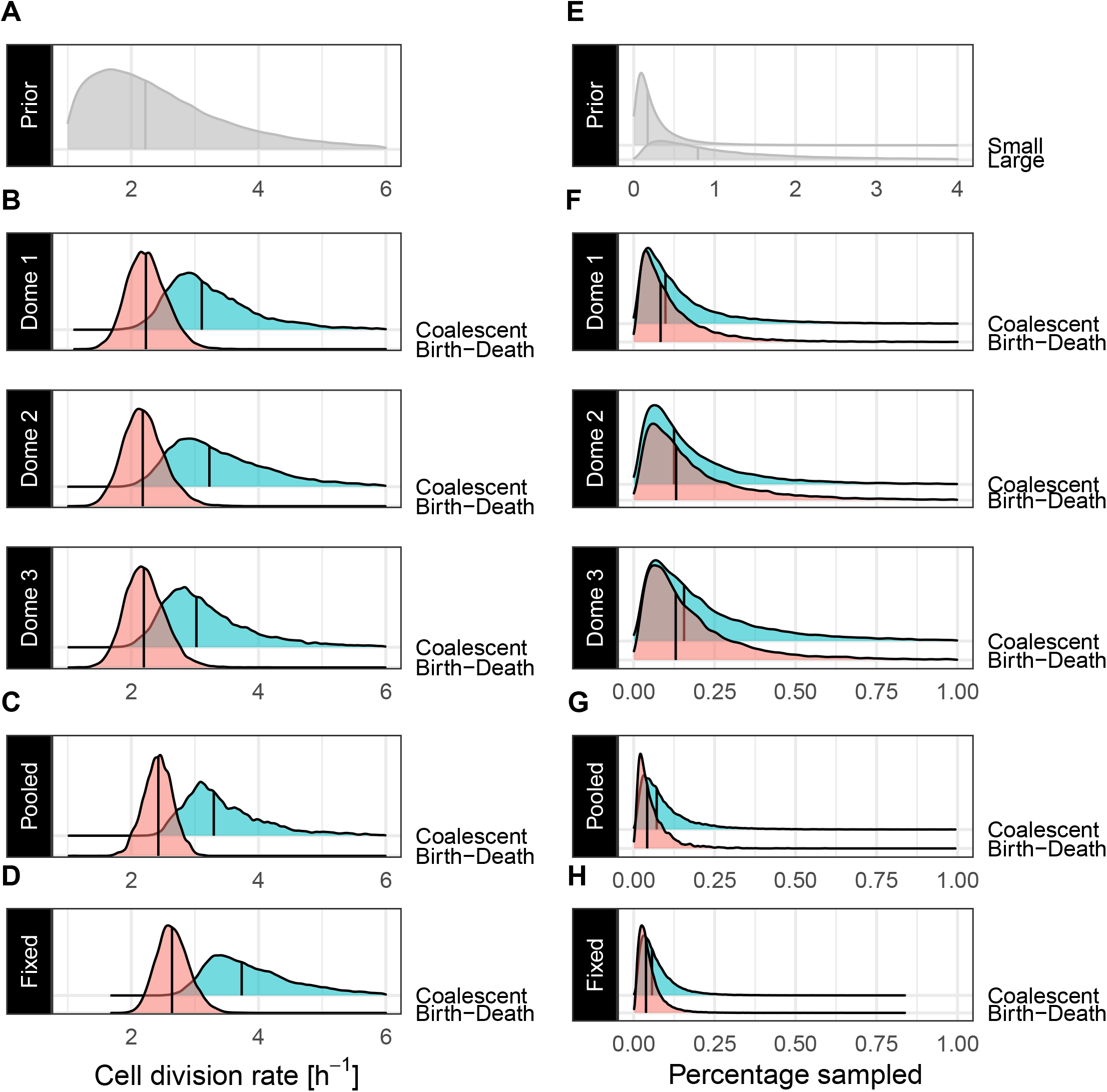
Phylodynamic inference of cell division rates (left) and sampling proportions (right) for dome stage (4.33 hpf) zebrafish Danio Rerio using 3 inference pipelines. (A) and (E) are the prior distributions chosen for both parameters of interest. We refer to the sampling proportion priors for alignments containing 20 sequences as *small* and for the fixed tree analysis with 100 sequences as large. (B) to (D) and (F) to (H) show posterior distribution obtained for the same parameters. In (B) and (F), we show parameters inferred independently for individual replicates of the experiment, and on alignments containing 20 sequences. in (C) and (G), we show estimates on pooled data. (D) and (H) are estimates using a fixed topology with 100 GESTALT sequences for replicate 1 (dome 1) of the experiment. On all panels, the black, vertical lines, represent the median for each distribution.

Next, we validated correctness of the likelihood implementation within its Bayesian inference framework with a simulation-based calibration procedure. For all parameters, after ensuring convergence, consistency between the credibility level and the percentage of HPD intervals containing the true value is achieved, showing that the targeted posterior distribution is sampled by GABI (Fig. 2 C). Additionally, we further investigated inferential properties of the GABI framework at credibility level of 95%. All parameters show the expected coverage at this level (within the 95% CI of the binomial distribution with n=100 trials and *p* = 0.95, see Appendix Table 6). Further inferential properties (RMSE, bias, relative HPD interval width) of the inference results under these validation settings are also reported in Appendix Table 7. Notably, these results are achieved with data generated under the assumption of a relatively low molecular clock rate. Therefore we underline that the performance shown here represents a lower bound on the inferential performance of GABI as we studied its properties under regime with low signal/little data. Higher mutation rates, leading to up to 7 indels per barcode as observed in our example datasets, may provide more signal and improve GABI’s performance.

### Phylodynamic analysis of dome stage zebrafish Danio rerio data

Estimates of the cell division rate and sampling proportion from dome stage zebrafish GESTALT data under the 3 chosen analysis pipelines are shown in Figure 2. Each compares estimates obtained under a constant birth-death model and the coalescent with exponential growth. While both model the growth of the studied population of cells as being exponential, their key difference is that the birth-death model assumes stochastic population dynamics, while the growth in the coalescent model is assumed to be deterministic. We consider both models to be potentially relevant in the context of this data. Overarching features of early embryonic development are thought to be largely deterministic, such as synchronicity until the 10th division [54, 55]. However, many genetic and environmental factors, including here, experimental conditions at play in generating replicates (such as, e.g. growth conditions, handling and harvesting) are also likely to introduce some level of stochasticity in the growth of the cell populations under study [56]. As the comparative adequacy of these two models for the analysis of growing cell populations remains to be investigated (in the context of epidemiological data, see [57]), we consider estimates under each model here as equally capable of capturing dome stage growth.

Individual division rate estimates under the birth-death-sampling model are consistent across all three replicates of the experiment, with median estimates around 2.2 *hr*^−1^, corresponding to cells until dome stage dividing on average every 27 minutes, (Fig. 2, B). While overlapping, respective estimates of the division rate under the coalescent model suggest a faster division rate, with all three median division rates above 3.1 *hr*^−1^, corresponding to divisions occurring every 20 min or less average. Our analysis infers a consistent sampling proportion for these three datasets and across both growth models used, where the alignments are estimated to contain between 0.1% and 0.2% of total cells in the embryos (Fig. 2, F). Comparatively, median estimates of parameters of the GESTALT mutation process are estimated with slightly more variation across replicates than population parameters, but do not suggest significant differences. For example, median molecular clock rate estimates range between 0.06 and 0.08 indels per site per hour across dome embryos (Fig. 3, E). Likewise, while site-specific cut rates vary across replicates, sites 3, 8, and 10 are consistently inferred to be the least active sites. Together, these estimate show that the V7 barcode design used in this experiment carries reproducible properties across the three replicates (see Appendix Fig. 3).

**Figure 3:**
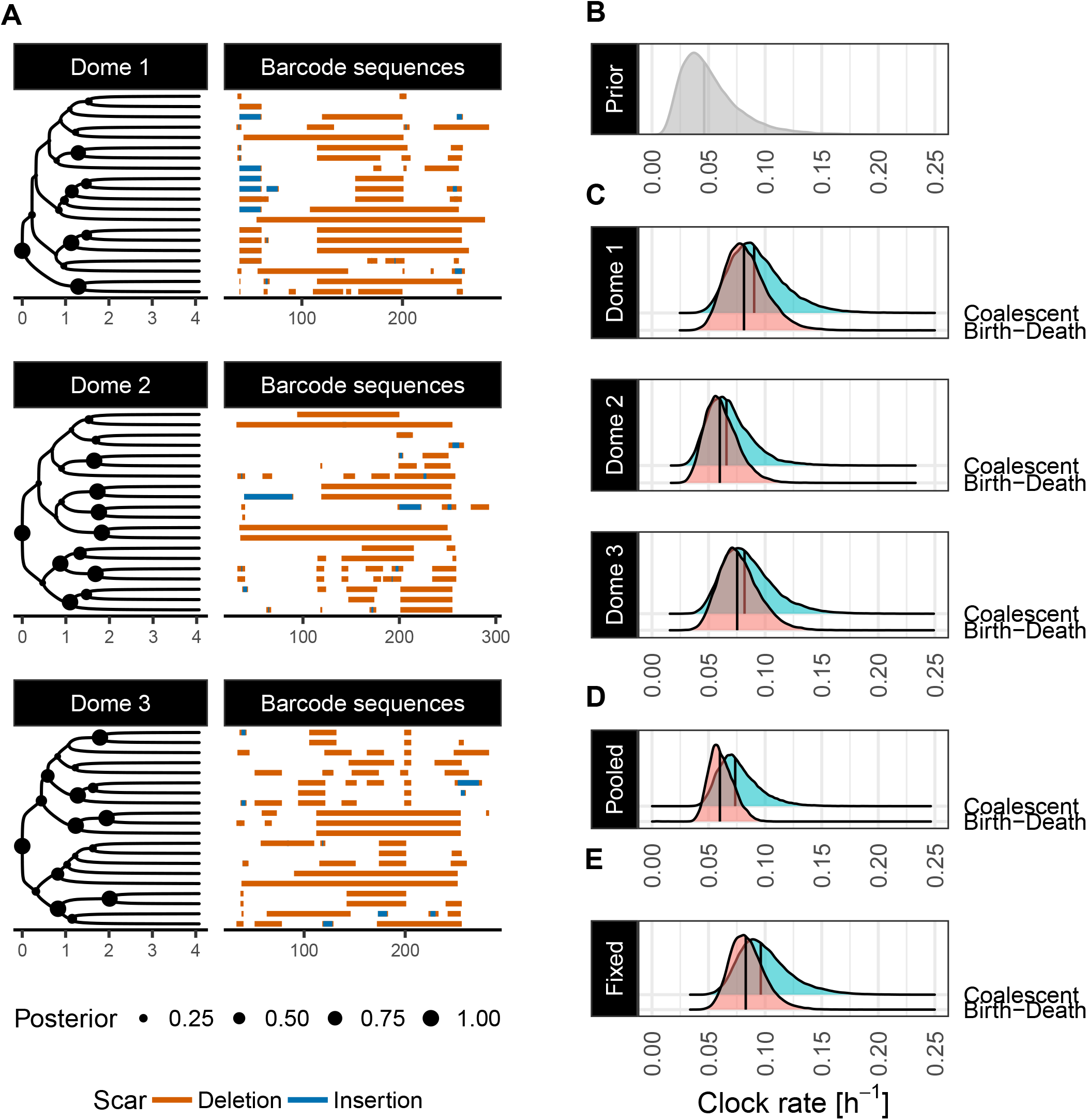
Phylogenetic inference for dome stage (4.33 hpf) zebrafish Danio Rerio. (A) Lineage trees inferred on individual replicates of the experiment, where nodes sizes represent posterior support. The time axis is in hours, and the axis corresponding to the barcode sequences represents the position (in base pairs) along the GESTALT barcode. (B) is the prior distribution chosen for the molecular clock rate, in number of indels inserted per site per hour, while (C) to (E) show its estimated posterior distribution under all 3 analysis pipelines.

We also examine the shape of the tree topologies inferred for the three datasets (Fig. 3 A), looking at the tree imbalance, which refers to the degree to which clade sizes, or groups of related cells, differ in the lineage tree. For a tree of size 20 under a birth-death-model, the expected imbalance index [58], representing the sum of the differences in clade sizes across all internal nodes, is 38.5, (95% CI [36.4, 41.1]). Interestingly, while the MCC tree for replicate 1 is within this expected range (40), for replicates 2 and 3, the inferred MCC lineage trees have lower imbalance indices (respectively, 13 and 20). The MCC trees show a similar pattern, where the second and third replicate obtained for the coalescent analysis are more balanced than the first replicate, with indices of 38 and 23 (51 for the first replicate). Under the assumption of uniform sampling of the cell population, more balanced trees correspond to an underlying process where subpopulations of cells are relatively even. This may agree with some of the assumptions of the population process under study: zebrafish embryos do not experience cell death or apoptosis until later embryonic stages and early cell divisions characteristic of zebrafish development are known to be largely synchronous until at least 3 hpf [55]. Based on this knowledge, highly balanced trees are expected to be reconstructed from such an experiment. While neither the birth-death-sampling nor the coalescent models specifically allow for synchronous divisions, the lower imbalance observed in the tree estimates for the second and third replicate may hint towards these alignments containing signal for such division events.

Our second analysis pipeline showcases how experimental replicates as analysed in our first example can be leveraged within a joint inference framework where cell division rates and mutation parameters are assumed to be the same across replicates (described in Fig. 1, C). In BEAST 2, this is easily achieved by multiplying the individual likelihoods for each replicate in each step of the MCMC. In Fig.2 (C and G) we show the division rate estimated jointly over all 3 datasets, under the birth-death and coalescent models. For the birth-death and coalescent models alike, the cell division rate inferred over this pooled data overlaps with individual estimates, but are inferred with more confidence. For example, the estimated cell division rate from pooled data under the birth-death model shows a a median 95%-level HPD interval width 26% narrower than the individual analyses. Pooled estimates under the exponential growth coalescent showcase a similar behavior, where the division rate is estimated with lower uncertainty (10% narrower 95%-level HPD interval width). Interestingly, both analyses suggest slightly elevated division rates when compared to individual replicates, with median cell division rates elevated on average by 0.18 *hr*^−1^ (coalescent) and 0.19 *hr*^−1^ (birth-death), and suggest lower sampling proportions (with sampling proportions of 0.06% and 0.07% for the birth-death and the coalescent, respectively).

As a third example of an analysis pipeline using GABI for phylodynamic inference, we show parameter estimates obtained using as input a fixed GAPML tree topology for the first replicate of the experiment (dome 1) (Fig. 2, D, H). In this scenario, only branch lengths are allowed to vary, where GABI provides estimates of the timing of individual cell divisions. This is consistent with many pipelines for phylodynamic inference in both fields of macroevolution and epidemiology (e.g.[59, 60]), where pre-existing, or separately inferred, fixed tree topologies are used as input to infer population dynamic parameters. This is particularly relevant in the context of Bayesian inference, where the exploration of the exponentially-growing lineage tree space quickly becomes intractable. Thus, using trees reconstructed with other, heuristic-based methods is often considered to enable the analysis of larger datasets. Here, we use a maximum-likelihood tree estimated under GAPML, for an alignment of 100 sequences as input, which is faster than GABI using their parsimony-guided tree search. The estimated division rate and sampling proportion under this pipeline are shown in Figure 2, D, H. Under the birth-death model, the division rate inferred remains mostly consistent with both previous analyses, but akin to the pooled analysis may suggest a slightly faster rate, 2.6 *hr*^−1^, with divisions occurring on average every 23 min. The division rate estimated for this dataset under the coalescent model is most shifted towards faster divisions, suggesting a median time until division of 16 min.

All three pipelines recover division rates that broadly agree with observed cell division lengths in zebrafish embryos: leading up to the mid-blastula transition (10th cycle post-fertilization), embryos undergo synchronous cell divisions every 18min a most (from 6 min in early cell divisions to between 12.9 to 18 min starting from the 6th division,[55]). After that stage, synchronicity is lost as individual cell cycles lengthen, and are found to last for up to an hour. While we do not model a time-varying rate for the purpose of this analysis, our estimates, considering both posterior medians and their uncertainty, successfully capture these known average dynamics.

## Discussion

In this paper, we described GABI, an implementation of an editing model and likelihood calculation in BEAST 2 enabling Bayesian phylodynamic inference using GESTALT data. Its first novel feature is that it allows to infer GESTALT cell lineage trees that are scaled in calendar time by incorporating a molecular clock model to the phylogenetic likelihood calculation. This feature makes GABI the only available method capable of inferring both the lineage topology and cell division times simultaneously using a mechanistic model of the GESTALT mutation process, where other methods [34, 61] require a fixed tree topology as input to generate chronograms. We validated this framework and presented three analysis pipelines to show its flexibility. These three pipelines are representative of most scenarios under which single-cell lineage tracing data may be analysed: GABI supports the pooled use of data from experimental replicates and enables both full phylogenetic inference or, alternatively, the use of fixed lineage topologies as input.

As such, GABI’s implementation enables the application of Bayesian phylodynamics to GESTALT datasets through the variety of tree priors readily available in BEAST 2. This opens the way for a vast range of studies investigating growth, apoptosis, and differentiation of cell populations through time. Thus, estimating birth and death parameters from trees, for which Yule 1925 [5] was the starting point, is here brought to the single cell application.

Our application of GABI to study the growth dynamics characteristic of dome stage zebrafish embryos is a first step showcasing these new inferential avenues using GESTALT barcode data. Our pipeline inferred cell division rates of zebrafish development by incorporating relevant experimental parameters as priors in our Bayesian inference, under the birth-death-sampling and the exponential growth coalescent model and across different analysis setups. This analysis showcases the ability for GESTALT data to help recover not only lineage relationships, but also capture signal for the growth patterns of cell populations.

While we showed a promising example of a first application, many other, more complex, datasets may require more work to accurately model certain features of both the mutation and growth processes. For example, one prominent feature of the GESTALT mutation process in zebrafish is tapering of the editing rate with time, which currently is not accounted for by the strict molecular clock model that we applied. Relaxed or time-dependent models may be more appropriate for analyses of later stage data [33, 46]. Like-wise, early cell divisions are known to occur with varying levels of synchronicity, and the cell cycle length is thought to lengthen in a time-dependent manner. Both of these properties are not captured by the models used in this study. Modelling the division process with readily available time-dependent models [7] and/or relaxing the assumption of exponentially distributed waiting times of cell divisions may also need to be explored in follow-up studies [62]. Notably, both birth-death and exponential growth coalescent models do not allow for synchronous binary cell divisions known to be characteristic of early vertebrate embryonic development. Future work may need to investigate the extent to which this biases estimates of the cell division rate. This motivates future work addressing the adequacy of these existing models to capture the growth dynamics specific to cell populations, and assess the need for new, custom models of cell proliferation.

GABI’s computational limitations lie in that the likelihood calculation becomes computationally prohibitively intensive for datasets containing more than hundreds of sequences. This remains a major challenge, as GESTALT alignments can contain in the range of thousands of sequences. The main contributor to this issue is the ancestral state pruning algorithm, which scales poorly in the presence of large inter-target deletions that potentially mask pre-existing indels. In [37], this issue is dealt with for large trees with imposing additional constraints on the number of editing event on each branch. Implementing this feature in GABI may be one way to speed computation. Additionally, GAPML only tunes topologies that preserve the maximum parsimony score. The combined use of such heuristics, or further approximations, may alleviate runtime issues but likely at the expense of accuracy. Phylodynamics has had a tremendous impact in the fields of macroevolution and genomic epidemiology. We demonstrated here that single cell barcoding data shows similar promises for the field of cell biology. Time-scaled trees obtained from lineage tracing data can provide estimates of cell division and apoptosis rates. Further, the increasing availability of datasets that combine lineage tracing with additional modalities, such as transcriptomic data [63], and the use of more complex models to accommodate this data, such as multi-type models [64] should further allow to infer the process of differentiation – describing with increasing levels of detail the drivers of multicellular growth into a functioning organism. Thus we envision that single-cell Bayesian phylodynamics, through implementations such as GABI, will have similar impact in the field of cell biology as was the case for macroevolution and epidemiology.

## Acknowledgments

This project has received funding from the European Research Council (ERC) under the European Union’s Horizon 2020 research and innovation programme grant agreement no. 101001077. M.M. was supported by a Postdoctoral Fellowship funded by Eidgenossische Technische Hochschule Zurich. The authors would like to thank Timothy G. Vaughan, Ugnė Stolz, Nicola Mulberry and Marcus Overwater for helpful discussions and feedback.

## Conflict of interest

The authors declare no conflicts of interest.

## Authors’ Contributions

A.Z.: conceptualization, data curation, formal analysis, investigation, methodology, project administration, software, validation, visualization, writing—original draft, writing—review and editing; S.S.: conceptualization, methodology, supervision, writing—review and editing; M.M.: conceptualization, methodology, supervision, writing—review and editing. T.S.: conceptualization, methodology, project administration, funding acquisition, resources, supervision, writing—review and editing.

## Data accessibility

The codebase used to run all inferences shown in this article is available at: https://github.com/azwaans/GABI. All code used to process data and analyse inference outputs can be found at: https://github.com/azwaans/GABIanalysis

## Notes

https://github.com/azwaans/GABI

https://github.com/azwaans/GABIanalysis

### Competing Interest Statement

The authors have declared no competing interest.

