## Appendix for "Bayesian phylodynamics of early vertebrate development in BEAST 2"

### Bayesian phylodynamics of early vertebrate development in BEAST 2
### Supplemental material

Antoine Zwaans *[1,2], Sophie Seidel[1,2], Marc Manceau[1,2], and Tanja Stadler†[1,2,+]

[1]Department of Biosystems Science and Engineering, ETH Zürich, Basel, Switzerland
[2]Swiss Institute of Bioinformatics, Lausanne, Switzerland
[+]Corresponding author

31 May 2024

*Contact
†Contact

### 1 Appendix A

#### 1.1 Tables

| Parameter | Symbol | Value |
|---|---|---|
| Birth rate | $\beta$ | 0.8 |
| Death rate | $\lambda$ | 0.2 |
| Sampling proportion | $\rho$ | 0.0006 |
| Origin or experiment duration | $o$ | 25 |

**Table 1:** Parameters used to simulate the phylogenetic trees under a birth-death-sampling model for the quantitative validation.

| Parameter | Symbol | Distribution | 95% HDI |
|---|---|---|---|
| Clock rate | $r$ | Log-normal ($\mu = -4, \sigma = 0.2$) | [0.01, 0.02] |
| Site Cut rates | $\lambda_{i,i\in[1,4]}$ | Log-normal ($\mu = 0, \sigma = 0.2$) | [0.6,1.48] |
| Double cut weight | $\omega$ | Log-normal ($\mu = -3.1, \sigma = 0.2$) | [0.03,0.0667] |
| Long deletion factor | $\gamma_{j,j\in[1,2]}$ | Beta ($\alpha = 5, \beta = 70$) | [0.022,0.133] |

**Table 2:** Distributions for the editing model parameters used to simulate GESTALT alignments and as priors in the quantitative validation.

| Parameter | Symbol | Distribution | 95% HDI |
|---|---|---|---|
| Clock rate | $r$ | Log-normal ($\mu = -3.076, \sigma = 0.5$) | [0.0173, 0.123] |
| Site Cut rates | $\lambda_{i,i\in[1,4]}$ | Log-normal ($\mu = 0, \sigma = 0.5$) | [0.375,2.66] |
| Double cut weight | $\omega$ | Log-normal ($\mu = -3.1, \sigma = 0.2$) | [0.03,0.0667] |
| Long deletion factor | $\gamma_{j,j\in[0,1]}$ | Beta ($\alpha = 2, \beta = 32$) | [0.0074,0.158] |

**Table 3:** Prior distributions for GESTALT parameters in the analysis of dome stage embryos

| Parameter | Symbol | Distribution | 95% HDI |
|---|---|---|---|
| Division rate | $\beta$ | Log-normal ($\mu = 0.770, \sigma = 0.5$) | [0.81, 5.75] |
| Cell death rate | $\delta$ | 0 | fixed |
| Sampling proportion - 20 tips | $\rho$ | Log-normal($\mu = -6.3630, \sigma = 1.0$) | [0.0002, 0.01] |
| Sampling proportion - 100 tips | $\rho$ | Log-normal($\mu = -6.3630, \sigma = 1.0$) | [0.001, 0.06] |
| Origin | o | 4.33hrs | fixed |

**Table 4:** Prior distributions for the birth-death-sampling parameters in the analysis of dome stage embryos

| Parameter | Symbol | Distribution | 95% HDI |
|---|---|---|---|
| Growth rate | $g$ | Log-normal ($\mu = 0.770, \sigma = 0.5$) | [0.81, 5.75] |
| Population size at present | $N(t)$ | Log-normal($\mu = -9.357, \sigma = 1.0$) | [1600, 82000] |
| Origin | o | 4.33hrs | fixed |

**Table 5:** Prior distributions for parameters of the exponential growth coalescent in the analysis of dome stage embryos

| Parameter | Symbol | Coverage [%] |
|---|---|---|
| Clock rate | r | 97 |
| Site 1 cut rate | $\lambda_1$ | 90 |
| Site 2 cut rate | $\lambda_2$ | 92 |
| Site 3 cut rate | $\lambda_3$ | 90 |
| Site 4 cut rate | $\lambda_4$ | 91 |
| Double cut weight | $\omega$ | 96 |
| Left long deletion factor | $\gamma_0$ | 90 |
| Right long deletion factor | $\gamma_1$ | 94 |

**Table 6:** Coverages at the 95% credibility level in the validation study (percentage of times that the true value is contained in the inferred 95% HPD). At this level, coverages should be contained within the 95% CI of a draw from the Binomial distribution with n=100 and p=0.95, i.e. [90,99].

| Parameter | Symbol | RMSE | Bias | HPD width |
|---|---|---|---|---|
| Clock rate | r | 0.1057 | -0.029 | 0.68 |
| Site 1 cut rate | $\lambda_1$ | 0.180 | 0.029 | 1.06 |
| Site 2 cut rate | $\lambda_2$ | 0.248 | 0.014 | 1.04 |
| Site 3 cut rate | $\lambda_3$ | 0.255 | 0.255 | 1.08 |
| Site 4 cut rate | $\lambda_4$ | 0.274 | 0.095 | 1.11 |
| Left long deletion factor | $\gamma_0$ | 0.37 | 0.14 | 1.66 |
| Right long deletion factor | $\gamma_1$ | 0.28 | 0.059 | 1.54 |
| Double cut weight | $\omega$ | 0.175 | 0.078 | 0.74 |

**Table 7:** Average Root Mean Square Error (RMSE), Bias and Relative HPD widths for all parameters of interest (95 % credibility level)

#### 1.2 Figures

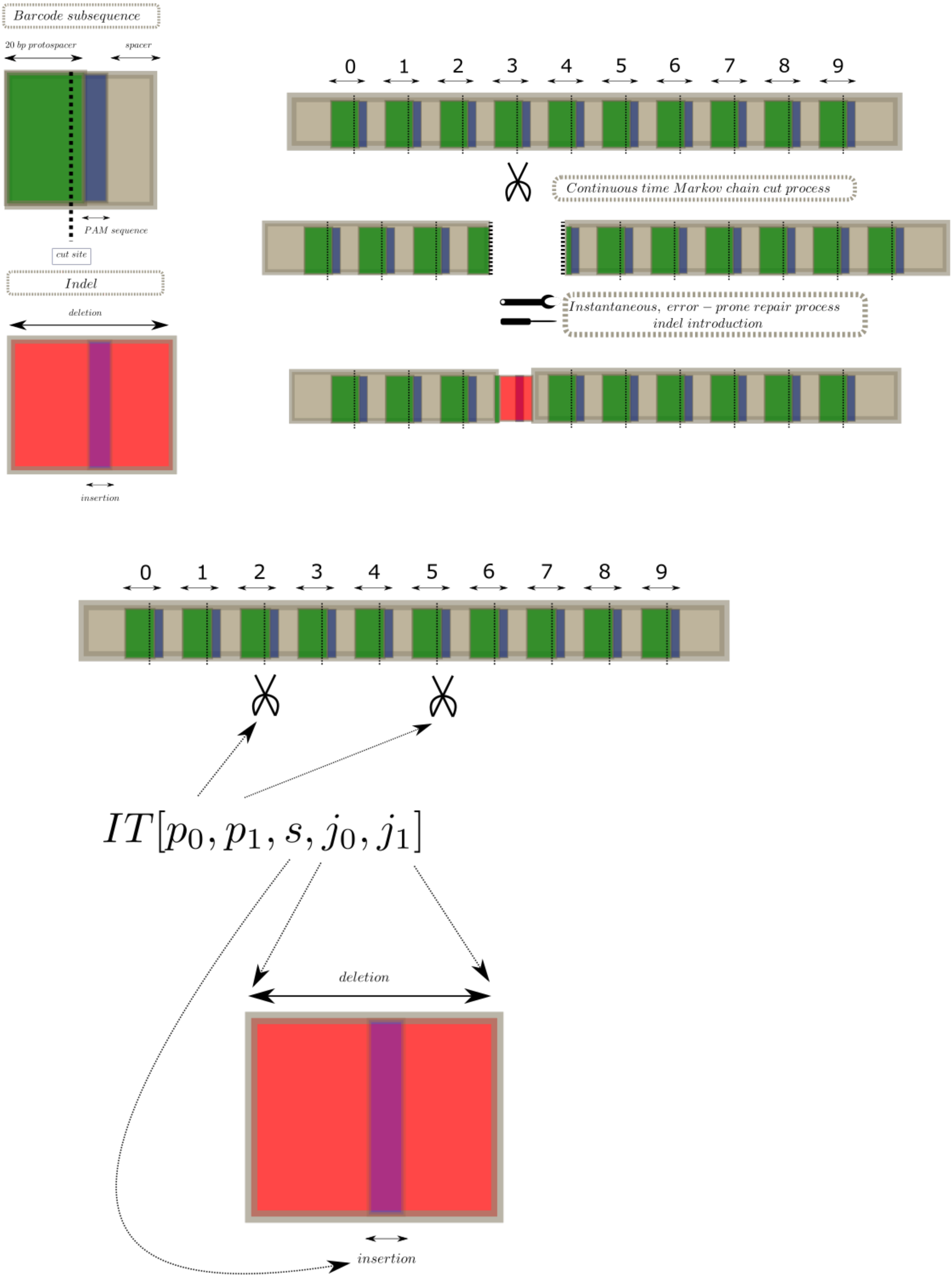


**Figure 1:** Depiction of the barcode mutation process and its building blocks. GAPML describes the GESTALT system with 2 building blocks (left): the arrayed target sequences (in green), and the indels introduced through the error-prone repair of double stranded breaks (in red). Indels blocks are introduced in the barcode as the result of the cutting and repairing processes occurring within target subsequences.

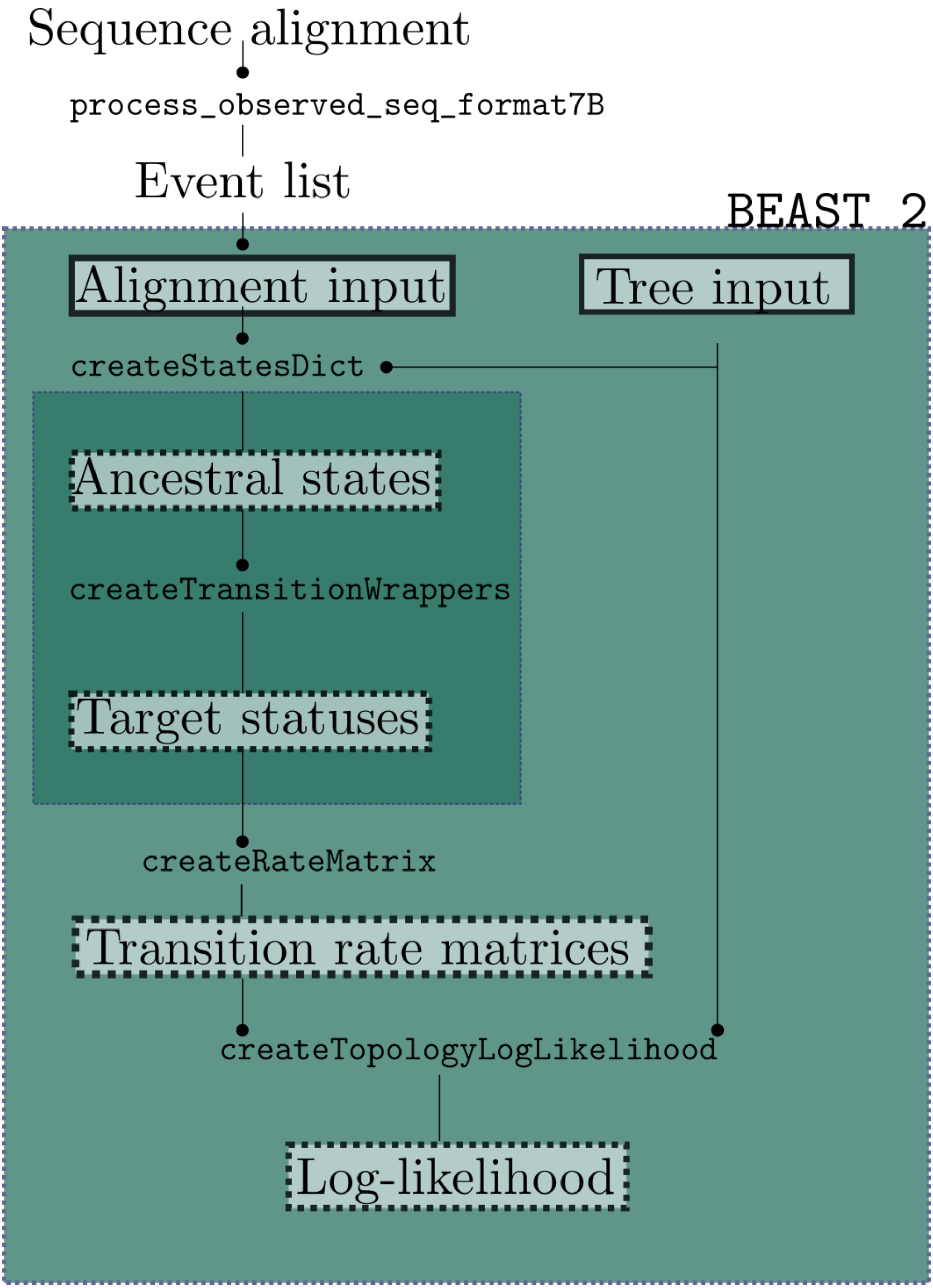


**Figure 2:** BEAST 2 Implementation architecture. Graphical representation of the GAPML objects and algorithms in BEAST 2. The arrow points represent inputs for each of the classes in the workflow.

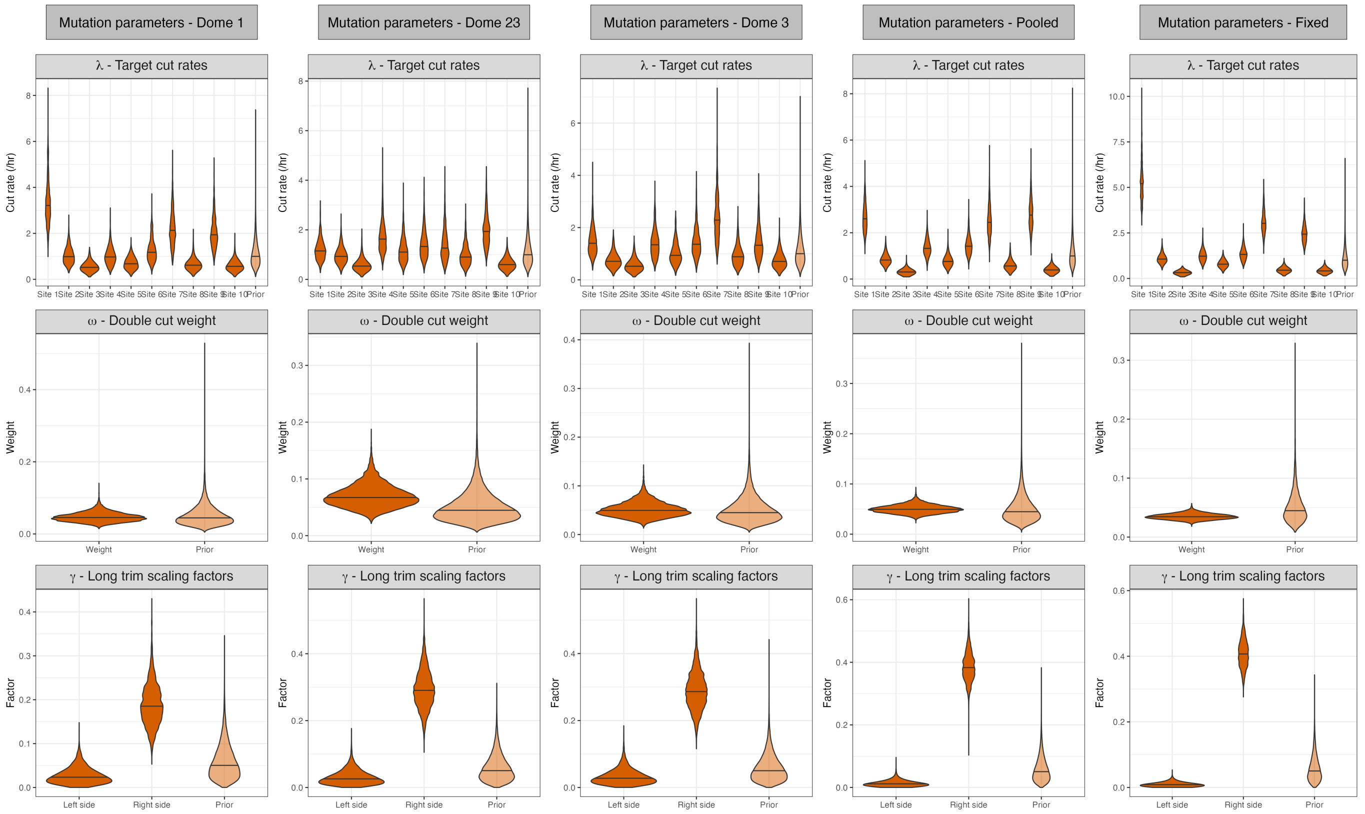


**Figure 3:** Parameters of the GAPML model estimated for each of 3 analysis pipelines described, under the birth-death model.

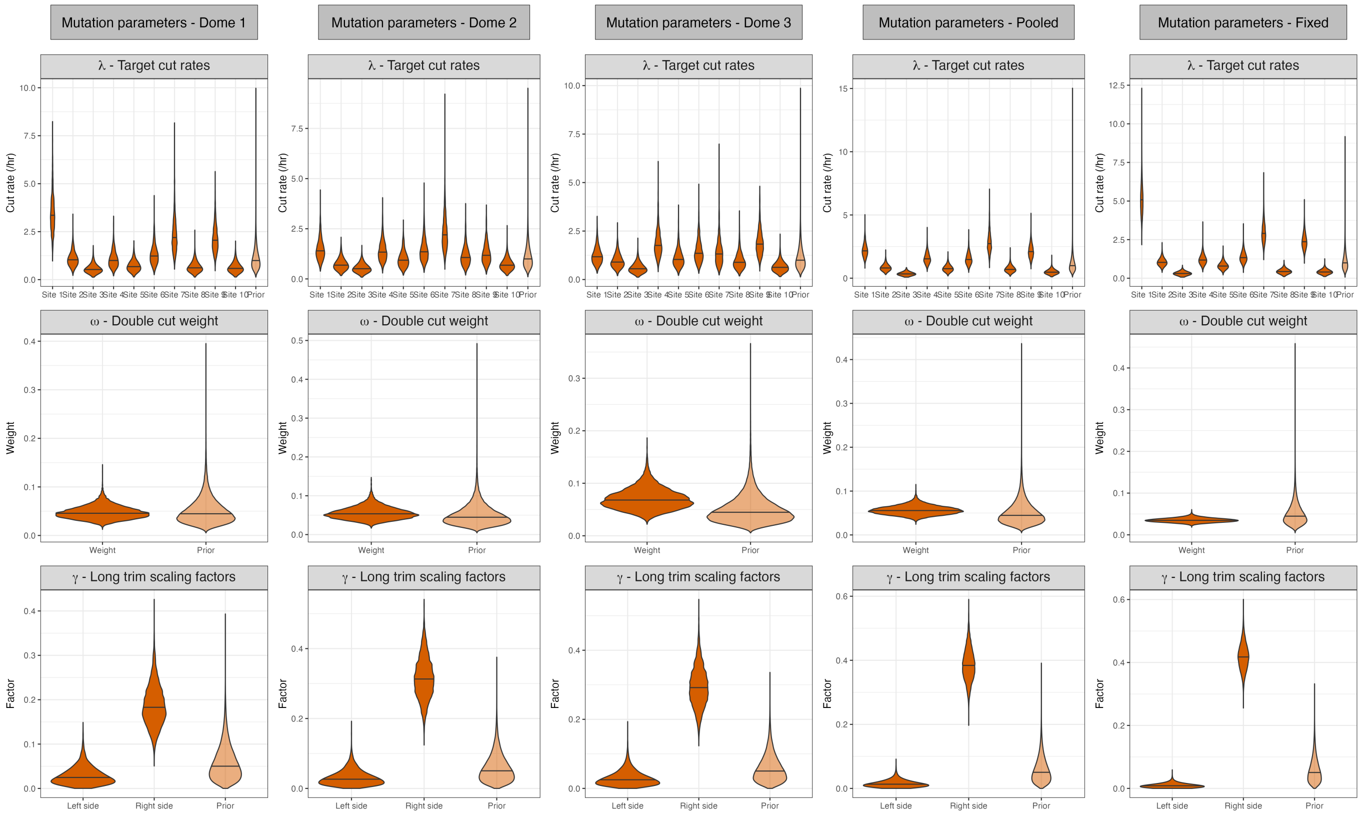


**Figure 4:** Parameters of the GAPML model estimated for each of 3 analysis pipelines described, under the exponential growth Coalescent.

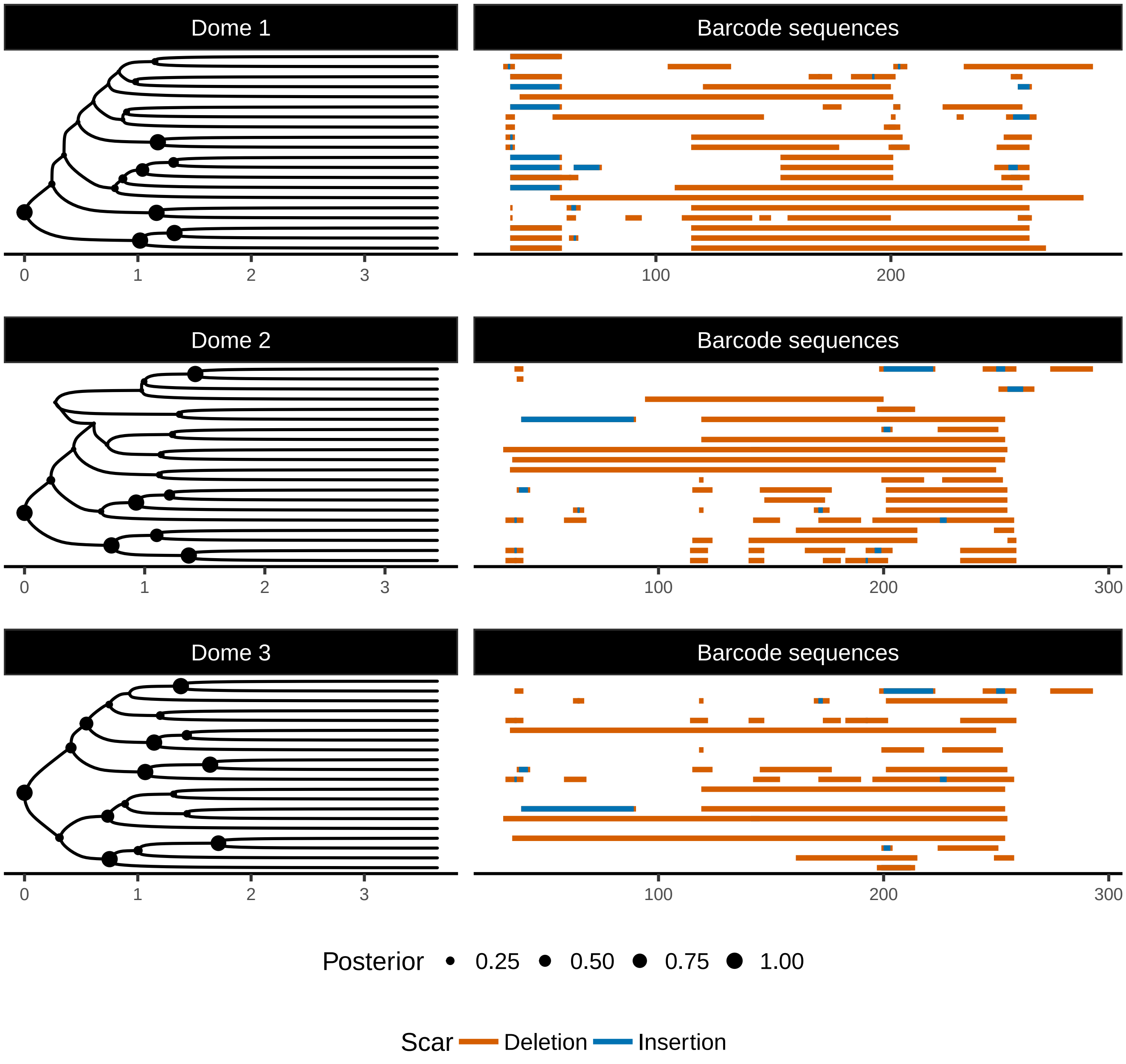


**Figure 5:** Lineage trees inferred on individual replicates of the experiment under the coalescent model, where nodes sizes represent posterior support. The time axis is in hours, and the axis corresponding to the barcode sequences represents the position (in base pairs) along the GESTALT barcode.

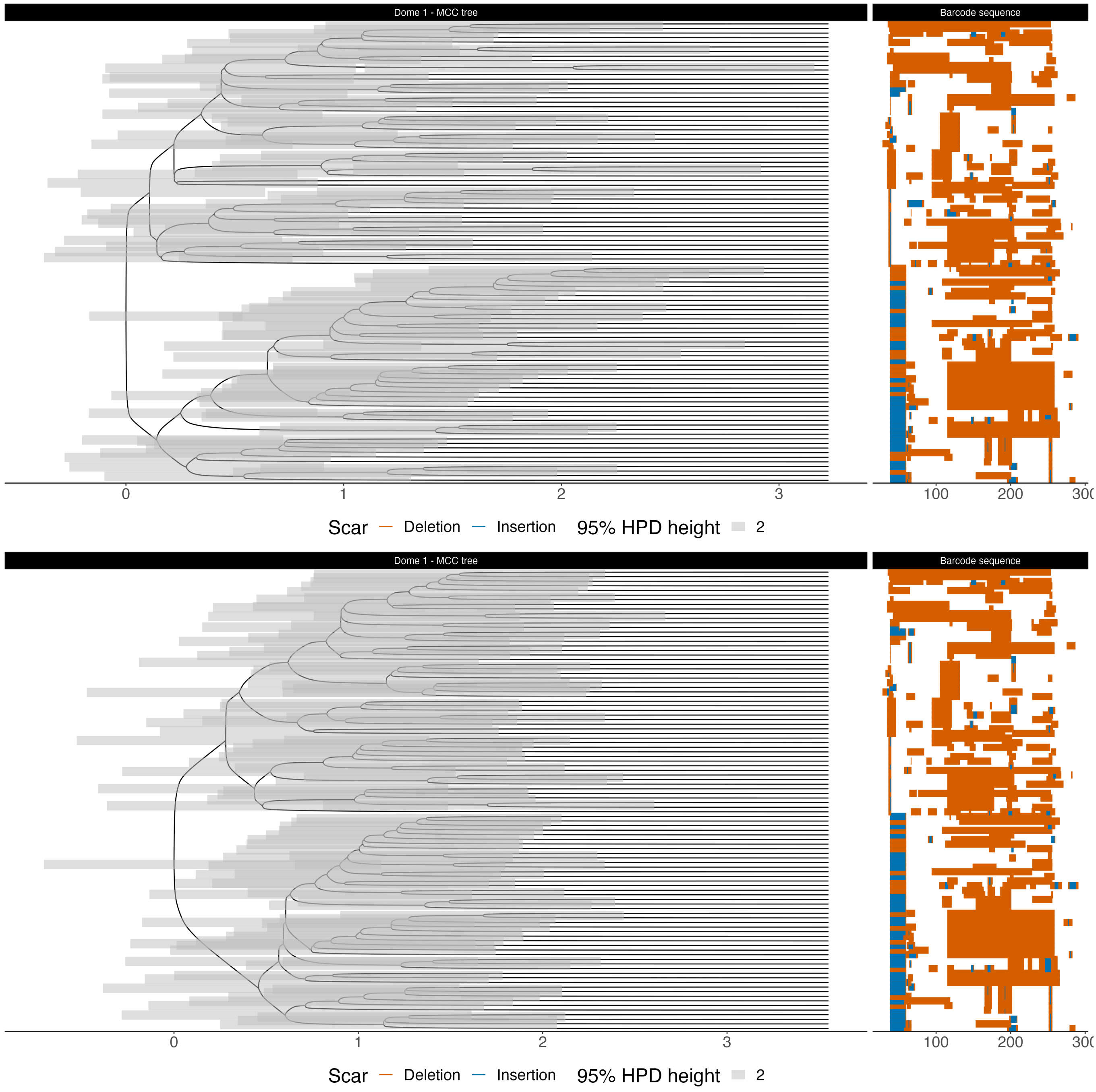


**Figure 6:** Time-scaled MCC, and uncertainty on divsion times (grey bars) tree estimated using GAPML, on a 100 sequence sample, corresponding to analysis pipeline 3, under the birth-death model, under the birth death model (top) and coalescent model (bottom). The axis ranges from the estimated root time to the present, while the origin of the process is fixed to 4.33 hours.

### 2 Appendix B

#### 2.1 GAPML - Allele evolution model

In GAPML, indels are assumed to be the result of at most 2 events of Cas9-mediated cuts at target sites in the GESTALT barcode. Based on this assumption, any indel introduced in a GESTALT barcode can be abstracted as an *Indel Tract*, denoted by $IT[p_0, p_1, s, j_0, j_1]$, where $j_0$ and $j_1$ ($j_0 \leq j_1$) denote which of its (at most 2) target sites were cut, $p_0$ and $p_1$ ($p_0 \leq p_1$) denote the absolute range of nucleotide positions deleted as a result, and $s$ denotes the sequence inserted (see Appendix Fig. 1, lower half). Using ITs as individual characters making up mutated GESTALT barcodes, if follows that single mutation steps consist of the introduction of an IT $d$ into a barcode sequence. This process is mathematically described as a continuous-time Markov chain (CTMC) on the space $\Omega$ of all such alleles in [1], where the mutation of a GESTALT barcode up to time $T$ is denoted as $\{X(t) : 0 \leq t \leq T\}$, and where each transition in state space is a discrete and irreversible mutation step that consists of applying an IT $d$ to the barcode sequence in state $a$, denoted as *Apply(a, d)*. Therefore, an allele is defined as a disjoint collection of at most $M$ indel tracts in the barcode:

$$a \equiv \{IT[p_{0,k}, p_{1,k}, s_k, j_{0,k}, j_{1,k}] : k \in \{1, M\}\} \tag{1}$$

#### 2.2 GAPML - Assumptions on the deletions

Deletions may extend to either sides of a cut, and, in the event of a double cut, where ($j_0 < j_1$), the entire range of positions between the 2 cuts is deleted, resulting in an *intertarget deletion*. The range of positions that can be deleted right and left of a cut is assumed to be bound by the neighboring sites' cut positions, i.e. an indel tract that cuts sites $j_0, j_1$ can only delete a range of positions that fulfills $c(j_0 - 1) < p_0 \leq c(j_0)$ and $c(j_1) \leq p_1 < c(j_1 + 1)$, where $c(j)$ denotes the absolute position of target cut site $j$ along the sequence. For a given indel spanning several targets, this means that the cut sites are always assumed to be the outermost positions. Deletions and insertion lengths are assumed to be Poisson distributed.

#### 2.3 GAPML - Partitioning of allele space $\Omega$

All alleles $a \in \Omega$ are defined by which of their sites are active, i.e. still amenable to cuts, and which are inactive, i.e. already harbour an indel. This is due to the introduction of an IT being irreversible as it results in the deactivation of the target sites $j_0, j_1$ it affects. This property maps alleles to a binary representation of the status of its barcode targets, denoted as $TargStat(a)$, where:

$$TargStat(a) = (TargStat(1; a), ..., TargStat(M, a)) \tag{2}$$

And where $TargStat(k; a)$ is the editing status of target $k$ in allele $a$ (0:active, 1:inactive) , such that:

$$TargStat(k; a) = 1\{\exists IT[p_0, p_1, s, j_0, j_1] \in \ a \text{ s.t. } j_0 \leq k \leq j_1\} \tag{3}$$

There are exactly $2^M$ of such target statuses. This property partitions the allele space $\Omega$ into groups of alleles that share a TargetStatus, which allows to study the behaviour of an aggregate, or "lumped" Markov process instead of the full Markov process ([2]). Transitions between two different target statuses happen as a result of any indel affecting the same set of targets. These supersets of indels are called *target tracts* (TT), and are defined by the range of targets they cut and deactivate. Analogue to the Markov chain describing the indel mutation process described above, single mutation steps in the "lumped" Markov process consist of applying TT to the allele. A target tract that cuts targets $j_0$ and $j_1$ and deactivates targets $j'_0$ and $j'_1$, is denoted by:

$$TT[j'_0, j_0, j_1, j'_1] = \{IT[p_0, p_1, s, j_0, j_1], \ p_0 \in pos(j'_0), \ p_1 \in pos(j'_1)\} \tag{4}$$

Where $pos(j'_0)$ and $pos(j'_1)$ denote the absolute nucleotide positions of cut sites contained in $j'_0$ and $j'_1$ within the barcode. Analogously to applying indel tracts to alleles in each mutation step, target tracts are successively applied to TargetStatuses. Thus, a set of target tracts induces a target status.

#### 2.4 Likelihood calculation

In molecular phylogenetics, the phylogenetic likelihood targets the probability of observing a sequence alignment given a lineage tree under a model of molecular evolution. Felsenstein's pruning algorithm [3] is used to calculate this likelihood efficiently by summing over all possible internal node states for a given tree using dynamic programming. For DNA sequences under evolutionary models of substitution, those states are exclusively one of the 4 nucleotides. In principle, the problem with using GESTALT barcodes as sequence data instead is that there are infinitely many possible indel states to sum over, which is computationally intractable. By considering the aggregate continuous-time Markov process described above instead of the full indel process, it was shown that the phylogenetic likelihood can be calculated by summing over at most $2^M$ allele statuses. To further avoid marginalizing over all of these $2^M$ states, GAPML introduces an algorithm used to prune which of these statuses are possible at each of the nodes in the lineage tree.

At each of the $2n-1$ nodes of a lineage tree $T$ with $n$ tips, the set of possible aggregated allele states at node $i$ $A_i$ that could have preceded the observed barcode sequences are computed using a recursive algorithm. These states are then used to target the probability density $P(D|T,\eta)$ of a GESTALT alignment given the tree and model parameters $\eta$, or phylogenetic likelihood.

#### 2.5 Transition probability matrix construction

Individual transition probabilities between any 2 states $a$ and $b$ of $\Omega$ under the model introduced above are obtained from a transition probability matrix constructed on each branch of the lineage tree to account for transitions between possible ancestral states as identified above. The rate at which an indel $d$ is edited into an allele $a$ $q(a, Apply(a,d))$, is factorised into 2 components, for each of the 2 steps of the mutation process: (1) the rate of applying a Target Tract $\tau = TT[j_0', j_0, j_1, j_1']$ into the allele, corresponding to the transition between aggregate states of the process $h(\tau, TargetStat(a))$, and (2) a probability scaling for the lengths of resulting insertions and deletions $Pr(d|\tau)$ (i.e. repairing the cut):

$$q(a, Apply(a,d)) = h(\tau, TargetStat(a))Pr(d|\tau) \tag{5}$$

All such transition rates in space $\Omega$ can be used to construct a transition rate matrix $Q$.

#### 2.6 GAPML - Parameterisation

The set $\eta$ of parameters of the GESTALT mutation model consists of the following. Each target site $k$ is considered to have its own, independent cut rate $\lambda_k$. Cuts that affect a single site at a time are distinguished from cuts that occur for 2 targets simultaneously, that result in *intertarget deletions*. These double cuts are assumed to occur at a rate that scales with the sum of the cut rates from the 2 sites it affects, multiplied with a weight $\omega$. The relative propensities at which the repair process results in long deletions, i.e., affecting neighboring sites, left and right of the cut are parameters $\gamma_0$ and $\gamma_1$ respectively. Combining these parameters, the rate $h(\tau, TargetStat(a))$ at which the cuts and deletions that result in a target tract $\tau = TT[j_0', j_0, j_1, j_1']$ are introduced into an allele $a$ with editing status $TargetStat(a)$ (0: is active, 1:inactive), are the following:

$$h(\tau, TargetStat(a)) = h_0(j_0, j_1, TargetStat(a))h_1(j_0, j_1, j_0', j_1') \tag{6}$$

Where the term $h_0$ specifies the rate at which the cuts are introduced:

$$h_0(j_0, j_1, TargetStat(a)) = \begin{cases} \omega(\lambda_{j_0} + \lambda_{j_1})1\{TargetStat(j_0, a) = 0\}1\{TargetStat(j_1, a) = 0\} & \text{if } j_0 \neq j_1 \\ \lambda_{j_0}1\{TargetStat(j_0, a) = 0\} & \text{otherwise} \end{cases} \tag{7}$$

And term $h_1$ scales that rate to consider the frequency of the deletions associated with that cut (affecting the neighboring site or not):

$$h_1(j_0, j_1, j_0', j_1') = \prod_{i=0}^{1} \gamma_i 1\{j_i \neq j_i'\} + 1\{j_i = j_i'\} \tag{8}$$

The lengths of insertions and deletions are all assumed to be Poisson distributed. Based on this property, the probability $Pr(d|\tau)$ for the lengths of deletions left and right of the cut sites and insertion length is obtained as the probability of these lengths under of their respective Poisson distributions in GAPML. The expressions for this factor are described in the supplement of ([1]).
